# Development of Carboxymethyl Cellulose (CMC)-Based Bio-composites as a Functional Substitute for Single-Use Plastics for Active Food Packaging Applications

**DOI:** 10.64898/2026.02.22.707343

**Authors:** Rogerson Anokye, Kwadwo Boakye Boadu, Kingsford Owusu Boateng

## Abstract

The production of petroleum-based plastics used for packaging has led to significant environmental challenges in both aquatic and terrestrial ecosystems. Consequently, there is a growing need to explore viable alternatives to the usage of these conventional plastics. This study investigates the utilization of cellulose powder for producing of biodegradable plastics as a more sustainable substitute for petroleum-based materials. Bioplastic films were formulated with varying glycerol contents ranging from 0.5ml - 2.0ml. The glycerol served as a plasticizer to improve the mechanical properties of the films, which were subsequently subjected to biodegradability and tensile strength tests. Biodegradability was evaluated through soil burial tests, which revealed that higher glycerol concentrations accelerated rate of weight loss, with the 2.0 ml formulation exhibiting the fastest degradation rate. Tensile strength increased with glycerol content up to 1.5 ml, where a maximum strength of 7.23 N/mm^2^ was recorded, but declined at 2.0 ml. The findings indicate that a glycerol concentration of 1.5 ml yields the most optimal bioplastic formulation for short-term packaging applications.

## 1.0 Introduction

Cellulose-based plastics have garnered significant attention as sustainable materials for applications including packaging, coatings, gels, foams, and composites (Zhang et al., 2022), Cellulose is a naturally abundant polymer sourced primarily from plants (George et al., 2021), algae (Nanda & Bharadvaja, 2022; Nandakumar et al., 2021), and bacteria (Meereboer et al., 2020), and is especially prevalent in wood, which contains 40–50% cellulose (Krumm et al., 2016). The extraction of cellulose involves processes such as alkalization, bleaching, and acid hydrolysis to yield micro- or nanocellulose (Magalhães et al., 2023; Wai & Ko, 2022). Historically, cellulose has been foundational in the development of early thermoplastics and continues to play a crucial role in manufacturing paper, rayon, and cellophane (Manali Shah et al., 2021).

Bioplastics, defined by their bio-based origin and biodegradability, decompose through microbial enzymatic activity and are increasingly utilized in packaging products with diverse shelf lives (Yaradoddi et al., 2020). Their production from renewable feedstocks, including underutilized waste streams like food residues and wood chips, offers a strategic reduction in reliance on fossil fuels (Kumar & Thakur, 2017). Biodegradability is governed by polymer structure rather than solely by feedstock origin (Pregi et al., 2023).

The bioplastics market encompasses both durable materials, such as bio-polyethylene, and degradable polymers including polybutylene succinate (PBS), polyhydroxyalkanoates (PHA), and polylactic acid (PLA) (Muhammad Shamsuddin, 2017). However, the integration of biodegradable plastics into conventional recycling streams presents challenges, complicating sorting processes and potentially diminishing the quality of recycled materials. Consequently, chemical and mechanical recycling methods are often preferred (Lamberti et al., 2020). It is noteworthy that not all bioplastics readily degrade in natural environments, and some fossil-based plastics may exhibit biodegradability depending on their molecular structure (Ezgi Bezirhan Arikan & Havva Duygu Ozsoy, 2015).

Despite increasing interest, bioplastics accounted for only 2% of global plastics production by 2018, with starch-based plastics and PLA dominating the market as of 2022. Meanwhile, petroleum-based plastic production continues to escalate, exacerbating environmental pollution (Rhodes, 2018). Global plastic production surged from 2 million tons in 1950 to 407 million tons in 2015, with packaging responsible for nearly half of resin use, having the shortest lifespan and generating most of the plastic waste. In 2015 alone, an estimated 302 million tons of plastic waste were produced, 97% of which originated from packaging, with projections estimating production could reach 1,124 million tons by 2050 (Rhodes, 2018).

The accumulation of plastic waste in terrestrial and marine environments poses severe ecological, health, and economic risks. Approximately 8.8 million tons of plastic enter oceans annually, contributing to an estimated 86 million tons of marine debris by 2013 (Geyer et al., 2017). This pollution threatens wildlife through ingestion, entanglement, and habitat degradation, while microplastics disrupt food webs and may impact human health (Geyer et al., 2017).

Bioplastics present a promising sustainable alternative capable of reducing long-term environmental persistence and supporting conservation efforts (Narancic et al., 2020). The growing global awareness and regulatory pressures are driving demand for renewable, biodegradable materials aligned with circular economy principles (Rosenboom et al., 2022). Biodegradable cellulose-based plastics decompose into benign compounds that can enhance soil quality and mitigate pollution across terrestrial, freshwater, and forest ecosystems (Adhikari et al., 2016).

Advancing the development and utilization of cellulose-derived bioplastics, such as those incorporating carboxymethyl cellulose (CMC), is therefore critical for addressing plastic pollution and fostering ecosystem resilience in the long term.

## 2.0 MATERIALS AND METHODS

### 2.1 Materials and equipment

The materials used in the study included carboxymethyl cellulose (CMC) powder (analytical grade, sodium salt form, medium viscosity), CMC served as the film-forming biopolymer. Glycerol (lab grade, ≥ 99% purity, liquid form) was used as a plasticizer to improve film flexibility. Vinegar (commercial food grade acetic acid solution, containing approximately 5% v/v acetic acid) used as mild acidifying agent to improve homogeneity of solution, and distilled water was used for all formulations to prevent interference with dissolved salts and impurities. Olive oil was smeared on the surface of plastic mould for easy removal of bioplastics.

The materials were obtained at Daniemma Medichem Ventures at Santasi, Kumasi. All reagents were used without further purification. The tensile strength tests were carried out at the Forestry Research Institute of Ghana (FORIG-Fumesua) and biodegradability tests were carried out at the Faculty of Renewable Natural Resources.

The equipment employed comprised a magnetic stirrer with integrated hot plate, borosilicate glass beakers, measuring cylinders, analytical balance (±0.001 g accuracy) spatula, plastic rectangular casting moulds and a Universal Testing Machine (Instron 3400/6800 series).

### 2.2 Method

#### 2.2.1 Preparation of bioplastics

Bioplastic samples were made at the General Chemistry Lab (Faculty of Renewable Natural Resources). A total of thirty-two (32) bioplastic film samples were prepared for this study. Sixteen samples (16) samples were designated for biodegradability testing, while the remaining sixteen samples were used for tensile strength analysis. The samples were divided into four formulations based on the glycerol content labelled 0.5 ml, 1.0 ml, 1.5 ml and 2.0 ml and it was replicated four independent batches as indicated in Table 1.

**Table 1.** Thickness in (mm) of the bioplastics for the biodegradability and tensile strength tests.

|  | 0.5 ml | 1.0 ml | 1.5 ml | 2.0 ml |
| --- | --- | --- | --- | --- |
| Biodegradability tests | 0.19 | 0.23 | 0.31 | 0.32 |
| Tensile strength tests | 0.21 | 0.24 | 0.29 | 0.27 |

Each formulation consisted of 4g of carboxymethyl cellulose (CMC) dissolved in 100 ml of distilled water and 2 ml of vinegar.

CMC powder was gradually added to distilled water and stirred at 600 rpm on a magnetic stirrer while heating at 80 °C for 30 minutes until a homogeneous solution was obtained. Glycerol and vinegar were subsequently added, and the mixture was stirred for an additional 10 minutes to ensure uniform plasticization and dispersion.

Equal volumes of each prepared solution were poured into clean, plastic rectangular casting molds to ensure uniform film thickness. The films were dried in the sun during the day and placed under ambient laboratory conditions (25 ± 2 °C, relative humidity 50–60% and natural airflow during the night for 3 days. After drying, the films were carefully removed from the molds and conditioned prior to testing. The formulation of the bioplastic casting solutions was expressed in terms of weight percentage (wt%). The wet casting solutions consisted predominantly of water (>92 wt%), with CMC (≈3.7 wt%), vinegar (≈1.8 wt%), and varying glycerol contents (0.6–2.3 wt%). For mechanical analysis, dry-basis composition was also considered, where CMC constituted 47–60 wt% of the film matrix and glycerol ranged from 9–30 wt%, depending on formulation. The dimensions of the bioplastic samples for the biodegradability and tensile tests are 4cm by 4cm and 2cm by 15cm respectively with varying thickness as shown in Figure 1 a and b.

**Figure 1.**
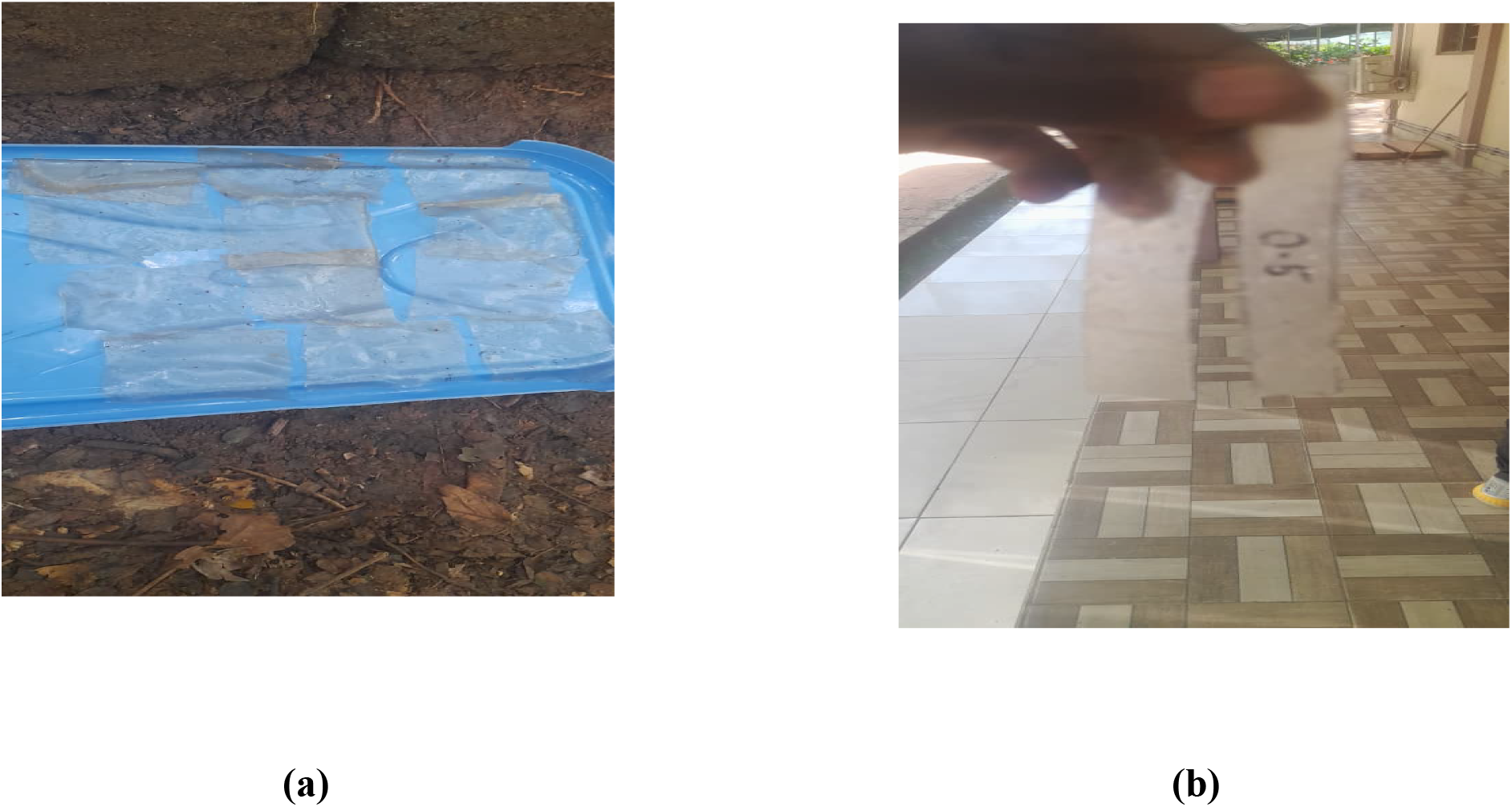
(a) Samples prepared for biodegradability test (b) Samples prepared for tensile strength tests.

All preparations were conducted using appropriate protective equipment (lab coat, gloves, and safety goggles), and care was taken when handling heated solutions and vinegar to prevent burns and splashes. The Universal Testing Machine was operated according to standard laboratory safety procedures to minimize mechanical hazards.

#### 2.3.1. Biodegradability Testing

Biodegradability of the bioplastic samples was assessed using a soil burial method in accordance with ASTM D5988-18. Samples measuring 4 cm by 4 cm were buried at a depth of 10 cm in a controlled compost soil environment maintained at 25°C and 50% relative humidity. The soil was regularly moistened to maintain consistent moisture content. Samples were retrieved at intervals of 24, 48 and 72 hours, gently cleaned of soil residues, dried at room temperature for an hour, and weighed to determine the weight loss percentage as an indicator of biodegradation as in Figure 2. Each test was performed in quadruplicate to ensure statistical reliability.

**Figure 2.**
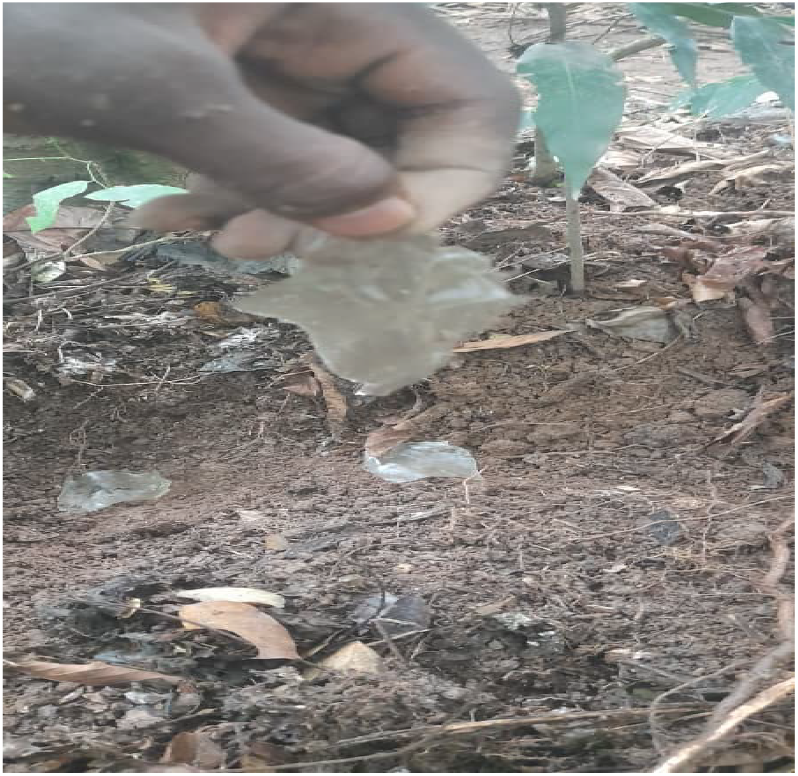
Samples being buried in the soil.

The percentage value of average mass reduction from buried bioplastics is obtained through the following equation:

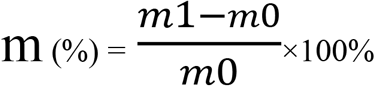

where m (%) is the percentage mass loss of the bioplastic sample m_1_ is the initial mass of the bioplastic sample before degradation m_0_ is the final mass of the bioplastic sample after degradation

#### 2.3.2 Tensile Strength Testing

Tensile strength was measured following ASTM D638 Type V standard using a universal testing machine (model and manufacturer) shown in Figure 3. Conditioned samples (ASTM D618) with dimensions conforming to Type V specimens were mounted and stretched at a crosshead speed of 5 mm/min until failure. The maximum load and elongation at break were recorded, and tensile strength and Young’s modulus were calculated. Four replicates per sample group were tested, and average values with standard deviations were reported.

**Figure 3.**
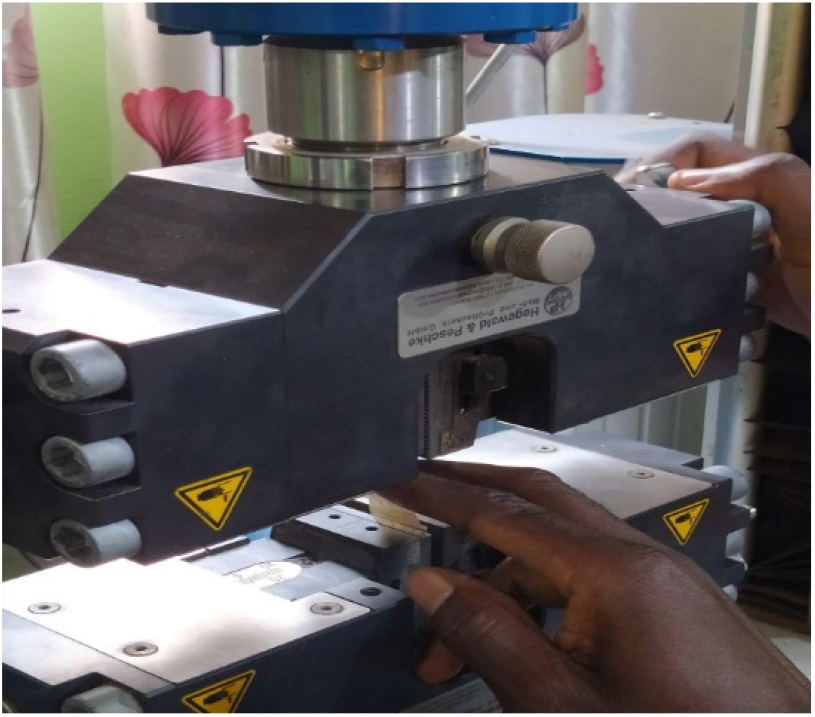
Samples placed in Universal Testing Machine.

## 3.0 RESULTS AND DISCUSSION

### 3.0 Tensile strength

The tensile strength of the CMC bioplastics was determined using a Universal Testing machine following. Formulations from glycerol concentration of 0.5ml, 1.0ml, 1.5ml and 2.0ml were prepared for the tensile testing as shown in Table 2. The stress at maximum load was recorded for all samples during tensile strength tests and the average was found across the four replicates The average tensile strength of bioplastic samples was evaluated across varying glycerol concentrations. Results demonstrated a progressive increase in tensile strength from 0.5 ml to 1.5 ml glycerol, with the highest tensile strength observed at 1.5 ml as shown in Table 2. However, a decline in tensile strength was noted at 2.0 ml glycerol concentration. The sample containing 0.5 ml glycerol exhibited the lowest tensile strength, which is attributed to insufficient plasticization leading to reduced polymer flexibility and cohesion. Consequently, this sample was more prone to premature fracture compared to those with higher glycerol content. The maximum tensile strength recorded was approximately 7.23 N/mm^2^ for the 1.5 ml glycerol sample, whereas the 2.0 ml glycerol sample showed a reduced tensile strength of about 6.09 N/mm^2^. This trend can be explained by the role of glycerol as a plasticizer (Dianursanti et al., 2018). Increasing glycerol content up to 1.5 ml enhanced polymer chain mobility and flexibility, facilitating improved stress distribution and resulting in increased tensile strength (Sapei et al., 2015). Beyond this concentration, excess glycerol disrupted intermolecular hydrogen bonding within the cellulose matrix, weakening polymer cohesion and thereby reducing tensile strength. These findings align with previous studies on bioplastics, where excessive glycerol similarly decreases tensile strength due to diminished crystallinity and structural integrity of the polymer matrix (Nasir & Othman, 2021).

**Table 2.**
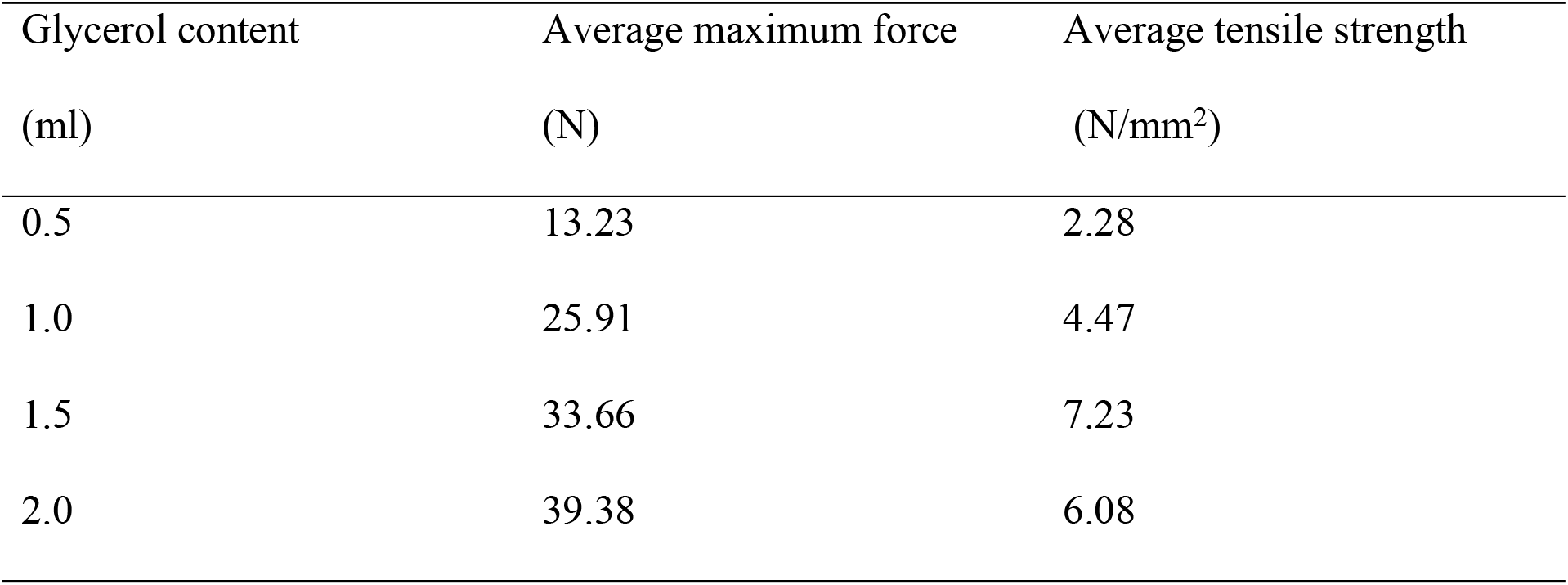
The average tensile strength and maximum force on various bioplastic samples.

| Glycerol content<br>(ml) | Average maximum force<br>(N) | Average tensile strength<br>(N/mm<sup>2</sup>) |
| --- | --- | --- |
| 0.5 | 13.23 | 2.28 |
| 1.0 | 25.91 | 4.47 |
| 1.5 | 33.66 | 7.23 |
| 2.0 | 39.38 | 6.08 |

### 3.2 Degradability of bioplastics

Biodegradability of the samples were assessed by soil burial tests according to the ASTM D5988 standards. The samples were buried at a depth of 10 cm below the surface of the soil and the weight loss of the bioplastics were observed at an interval of 24 hours for three days. Percentage of weight loss was used to indicate the rate of biodegradability of the bioplastics as indicated in Figure 5.

**Figure 5.**
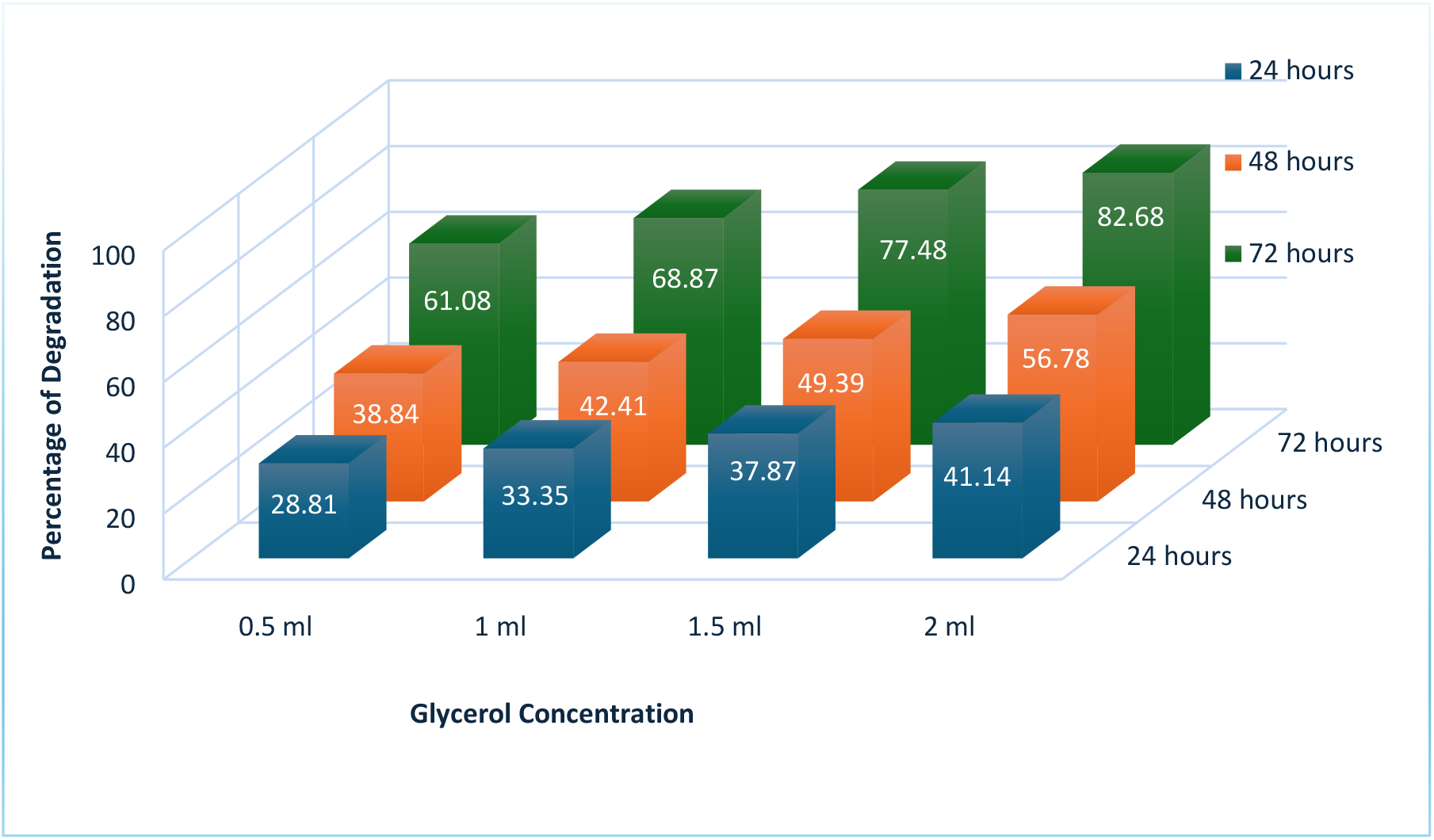
The average percentage weight loss for the different glycerol concentrations over the test period.

The results revealed a positive correlation between glycerol content and degradation rate as shown in all the three-time levels (Figure 5). Samples with lower glycerol content (0.5 ml) exhibited slower degradation when buried in soil for an equivalent duration compared to those with higher glycerol concentrations. In contrast, samples containing 2.0 ml glycerol demonstrated the most rapid degradation, indicating that increased glycerol content enhances the biodegradability of the bioplasticsas shown in Figure 6. This accelerated degradation at higher glycerol levels is attributed to glycerol’s capacity to increase water absorption and reduce the crystallinity of the bioplastic matrix (Bezirhan & Bilgen, 2019). These changes facilitate microbial colonization and enzymatic hydrolysis, thereby promoting faster breakdown of the material. These findings are consistent with previous studies on other bioplastics, where plasticizer concentration similarly influences degradation behavior (Fauziyah et al., 2021). The enhanced biodegradability of these bioplastics contributes to mitigating the environmental accumulation of plastics, supporting sustainability (Matsuura et al., 2008) and ecological health (Nandakumar et al., 2021).

**Figure 6.**
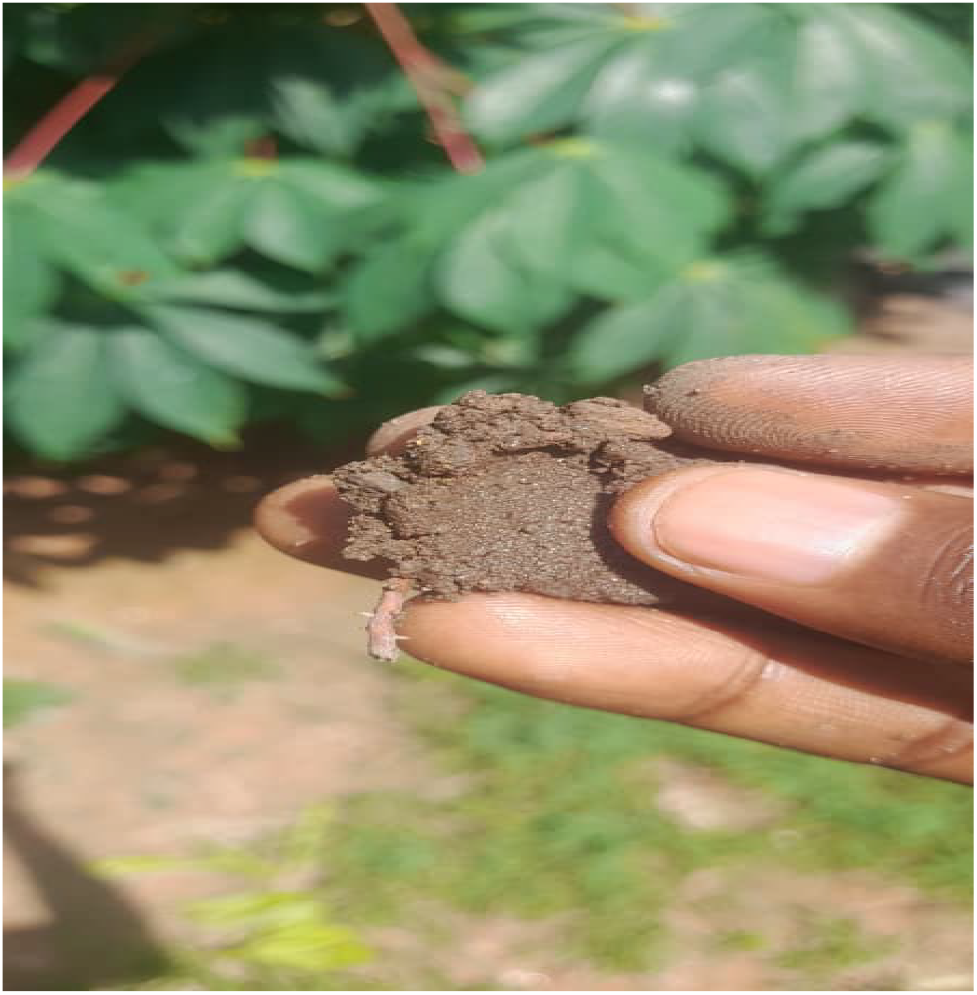
Bioplastic samples degraded after 72 hours.

## 4.0 CONCLUSION

The characterization of cellulose-based bioplastic composites reveals a non-linear relationship between plasticizer loading and material performance. Respirometric analysis conducted in accordance with ASTM D5988 demonstrates that increasing glycerol concentrations from 1.0 mL to 2.0 mL significantly accelerates microbial mineralization rates. Specifically, the 2.0 mL formulation exhibited the highest susceptibility to enzymatic cleavage, likely due to increased free volume within the polymer matrix, which facilitates moisture absorption and microbial infiltration.

However, the mechanical profile follows a divergent trajectory. Optimal tensile strength was localized at a 1.5 mL glycerol concentration, suggesting a threshold beyond which the plasticizing effect transitions into structural weakening via the disruption of intermolecular hydrogen bonding between cellulose chains. This observed trade-off confirms the strategic challenges identified by Matsuura, Ye, and He (2008): the optimization of bioplastics for commercial viability requires a precise calibration between “Design for Performance” (mechanical integrity) and “Design for Environment” (biodegradability).

Ultimately, while the 1.5 mL formulation offers a balanced mechanical-to-degradation ratio suitable for short-term packaging, the 2.0 mL variant is better positioned for applications requiring rapid organic recovery. These findings underscore the potential for cellulose-based substrates to serve as viable fossil-fuel mitigants, provided that end-of-life disposal infrastructure is aligned with the material’s specific degradation kinetics.

